# Discovery of a Novel Inhibitory Neuron Class, the L-Stellate Cells of the Cochlear Nucleus

**DOI:** 10.1101/2020.01.09.900092

**Authors:** Tenzin Ngodup, Gabriel E. Romero, Laurence O. Trussell

## Abstract

Auditory processing depends upon inhibitory signaling by interneurons, even at its earliest stages in the ventral cochlear nucleus (VCN). Remarkably, to date only a single subtype of inhibitory neuron has been documented in the VCN, a projection neuron termed the D-stellate cell. With the use of a transgenic mouse line, optical clearing and imaging techniques, combined with electrophysiological tools, we revealed a population of glycinergic cells in the VCN distinct from the D-stellate cell. These novel multipolar glycinergic cells were smaller in soma size and dendritic area, but over 10-fold more numerous than D-stellate cells. They were activated by auditory nerve fibers and T-stellate cells, and made local inhibitory synaptic contacts on principal cells of the VCN. Given their abundance, combined with their narrow dendritic fields and axonal projections, it is likely that these neurons, here termed L-stellate cells, play a significant role in frequency-specific processing of acoustic signals.

## Introduction

In the ventral cochlear nucleus (VCN), auditory nerve afferents make synapses onto multiple subtypes of excitatory neurons, setting up parallel streams of processing that are used by higher centers for processing auditory cues for sound intensity, frequency, and localization. This early-level activity of excitatory neurons is sculpted and stabilized by inhibitory neurons which use the transmitter glycine (Altschuler et al., 1986; Wenthold et al., 1986; Wu and Oertel, 1986; Kolston et al., 1992; Ferragamo et al., 1998a; Chanda and Xu-Friedman, 2010; Xie and Manis, 2013a, 2014; Manis et al., 2019). For example, glycinergic inputs onto excitatory bushy and T-stellate cells enhance acoustic tuning properties and temporal precision, and stabilize neuronal firing responses to acoustic inputs (Caspary et al., 1994; Kopp-Scheinpflug et al., 2002; Gai and Carney, 2008; Keine and Rubsamen, 2015). However, the sources of these inhibitory glycinergic inputs onto the principal cells in VCN are not well understood.

Within the VCN, only a single glycinergic inhibitory cell type, called the D-stellate or radiate multipolar cell, has been described (Oertel, 1983; Sento and Ryugo, 1989; Oertel et al., 1990; Doucet and Ryugo, 1997; Oertel et al., 2011; Xie and Manis, 2014). D-stellate cells are a small population of neurons (Campagnola and Manis, 2014) that send their axons to the ipsilateral dorsal cochlear nucleus (DCN) (Smith and Rhode, 1989; Oertel et al., 2011; Campagnola and Manis, 2014) but also to sites as distant as the contralateral cochlear nucleus (Schofield and Cant, 1996b; Needham and Paolini, 2003; Smith et al., 2005). Importantly, D-stellate cells provide broadband inhibition to their targets, as their large dendritic arbors receive input from a broad frequency spectrum of auditory nerve fibers (Smith and Rhode, 1989; Oertel et al., 1990). Another source of inhibition in VCN is a projection neuron originating in the DCN, the tuberculoventral cells (also called vertical cells) (Wickesberg and Oertel, 1988; Wickesberg et al., 1991; Xie and Manis, 2013b; Campagnola and Manis, 2014). Recent studies suggest that D-stellate and tuberculoventral cells do not fully account for the inhibition observed in VCN principal cells *in vivo* (Keine and Rubsamen, 2015) and *in vitro* (Campagnola and Manis, 2014). Indeed, no local inhibitory interneuron has ever been described for VCN, which is unlike most known brain regions. Early anatomical studies suggested the presence of short axon cells in the VCN (Lorente de Nó, 1981), or of inhibitory neurons different from the D-stellate cells, but the identity of such neurons remains unknown (Lorente de Nó, 1981; Wenthold, 1987; Doucet et al., 1999; Campagnola and Manis, 2014).

Here, we comprehensively examined the diversity of inhibitory neurons in the VCN using a well-characterized transgenic mouse line, GlyT2-GFP (Zeilhofer et al., 2005; Kuo et al., 2012; Moore and Trussell, 2017) which labels virtually all glycinergic neurons in the cochlear nucleus. With the use of this transgenic mouse, as well as optical tissue clearing, whole-cochlear nucleus-super-resolution microscopy, electrophysiological and morphological tools, we discovered a large population of inhibitory glycinergic cell types distinct from the D-stellate cell. The novel glycinergic neurons, which have a narrower receptive field than D-stellate cells, form the vast majority of inhibitory neurons in the VCN. We show that these cells, termed L-stellate cells, receive monosynaptic auditory fiber inputs and polysynaptic inputs from axon collaterals of T-stellate cells, and in turn locally inhibit bushy cells and T-stellate cells in the VCN. Thus, the VCN has a rich diversity of glycinergic interneurons to provide maximum flexibility to control the excitability of excitatory neurons in the VCN.

## Results

### Cell counts and soma size

We used a well-characterized GlyT2-GFP transgenic mouse (Zeilhofer et al., 2005; Albrecht et al., 2014; Moore and Trussell, 2017) in order to study the prevalence of glycinergic cells in the ventral cochlear nucleus (VCN). The neuronal glycine transporter, GlyT2, is a reliable marker of glycinergic cells and GFP is selectively expressed in > 90% of glycinergic cells in the cochlear nucleus (CN). To visualize all the glycinergic cells, we optically cleared whole CN (450-500 μm) using CUBIC-mount. Next, we imaged the whole CN, lateral to medial (Fig 1). Fig 1A shows a series of 50-μm thick image stacks of CN, lateral to medial. Not surprisingly, we observed a dense population of glycinergic cells in the dorsal cochlear nucleus (DCN) as described in previous studies (Oertel and Wu, 1989; Zhang and Oertel, 1993a, b; Kuo et al., 2012; Apostolides and Trussell, 2014). Also apparent were thick tracts of glycinergic fibers that entered the dorsal part of VCN, presumably projections of the glycinergic tuberculoventral neurons (vertical cells) in the DCN (Fig 1A). We also observed a lack of glycinergic cells in the octopus cell region of the posterior VCN, consistent with previous studies (Wickesberg and Oertel, 1988; Wickesberg et al., 1991). However, we found a large population of glycinergic cells distributed across the rest of the VCN. To obtain a global view of their distribution, the images were stitched and combined to create a 3D image of the entire CN (Fig 1B). The high density of glycinergic cells throughout VCN was surprising, because D-stellate cells are thought to be sparse, and the only other known glycinergic cells are Golgi cells, which are present mainly in the granule cell layer overlying VCN (Ferragamo et al., 1998b; Irie et al., 2006; Yaeger and Trussell, 2015).

**Figure 1.**
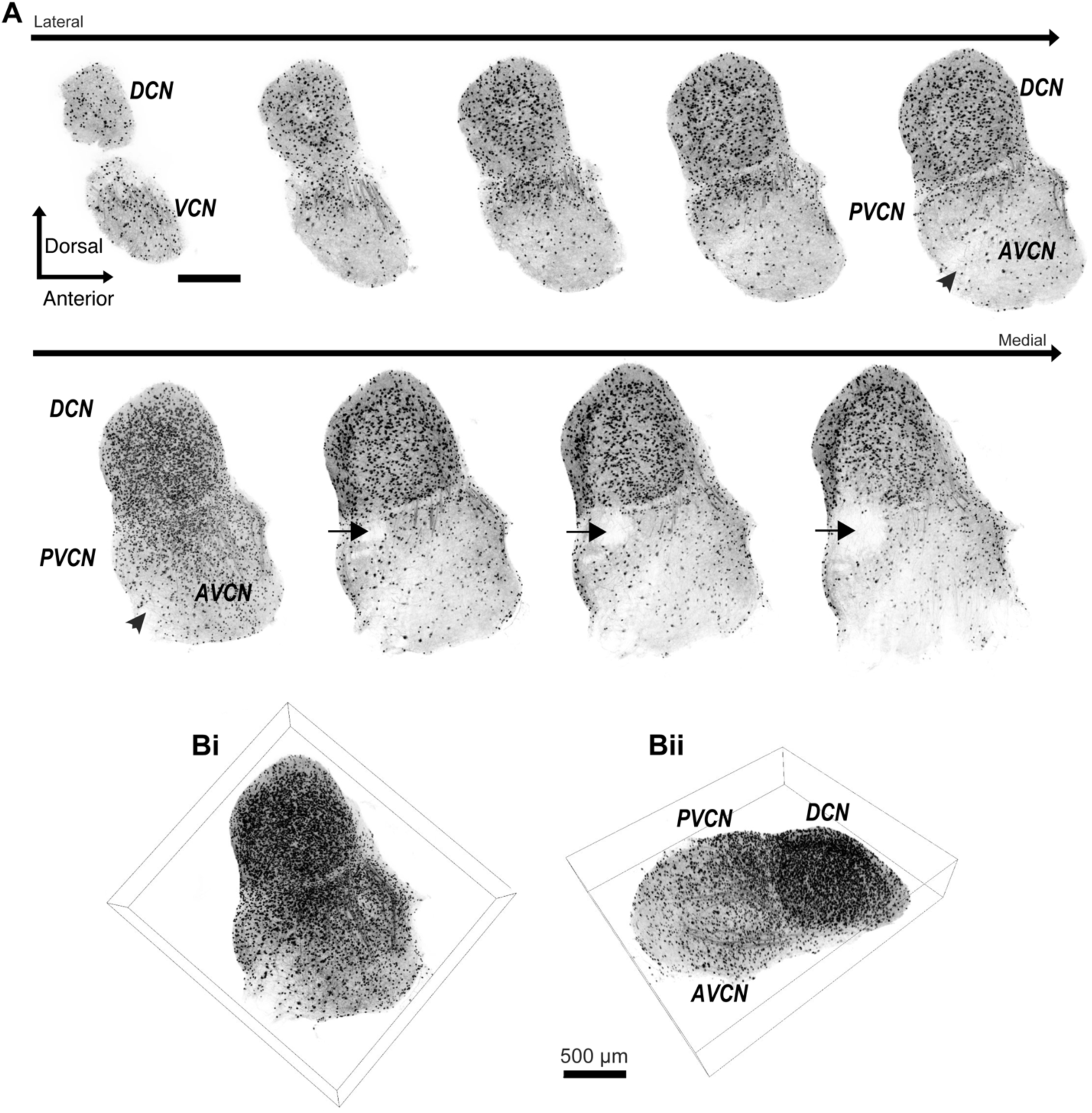
Diversity of glycinergic cell population in the CN. (**A**) Entire CN was optically cleared using CUBIC-mount and imaged with a super-resolution confocal microscope. Examples of 50-μm thick image stacks of CN from lateral to medial sides. DCN has a dense population of glycinergic cells. Octopus cell region in the PVCN (arrow) show lack of glycinergic cells. Arrowhead: auditory nerve root. (**B**) 3D images of entire CN show dense population of glycinergic cells not only the DCN but also in the VCN. Scale bar = 500 μm Figure 1-source data 1 The zip archive contains CN images used for the quantitative analyses shown in Figure 1 and 2. Images were collected at 5 μm steps using LSM 880 super resolution Airy scan micrcoscope with 25X objective

We restricted our analysis to neurons only in VCN by masking the area outside the VCN and then quantifying the glycinergic cell count using semi-automated ‘spot function’ in Imaris software (Fig 2A, B). This counting procedure yielded a total of 2706 ± 107 glycinergic neurons in the VCN (*n* = 4 VCNs, 3 mice). We next quantified the soma volumes of all glycinergic neurons using the ‘surface function’ in Imaris (Fig 2C, D), and found that the soma volume distribution was positively skewed (Fig 2E), such that the vast majority of the glycinergic cells had small somas, and a minority had large somas (Fig 2E, inset).

**Figure 2.**
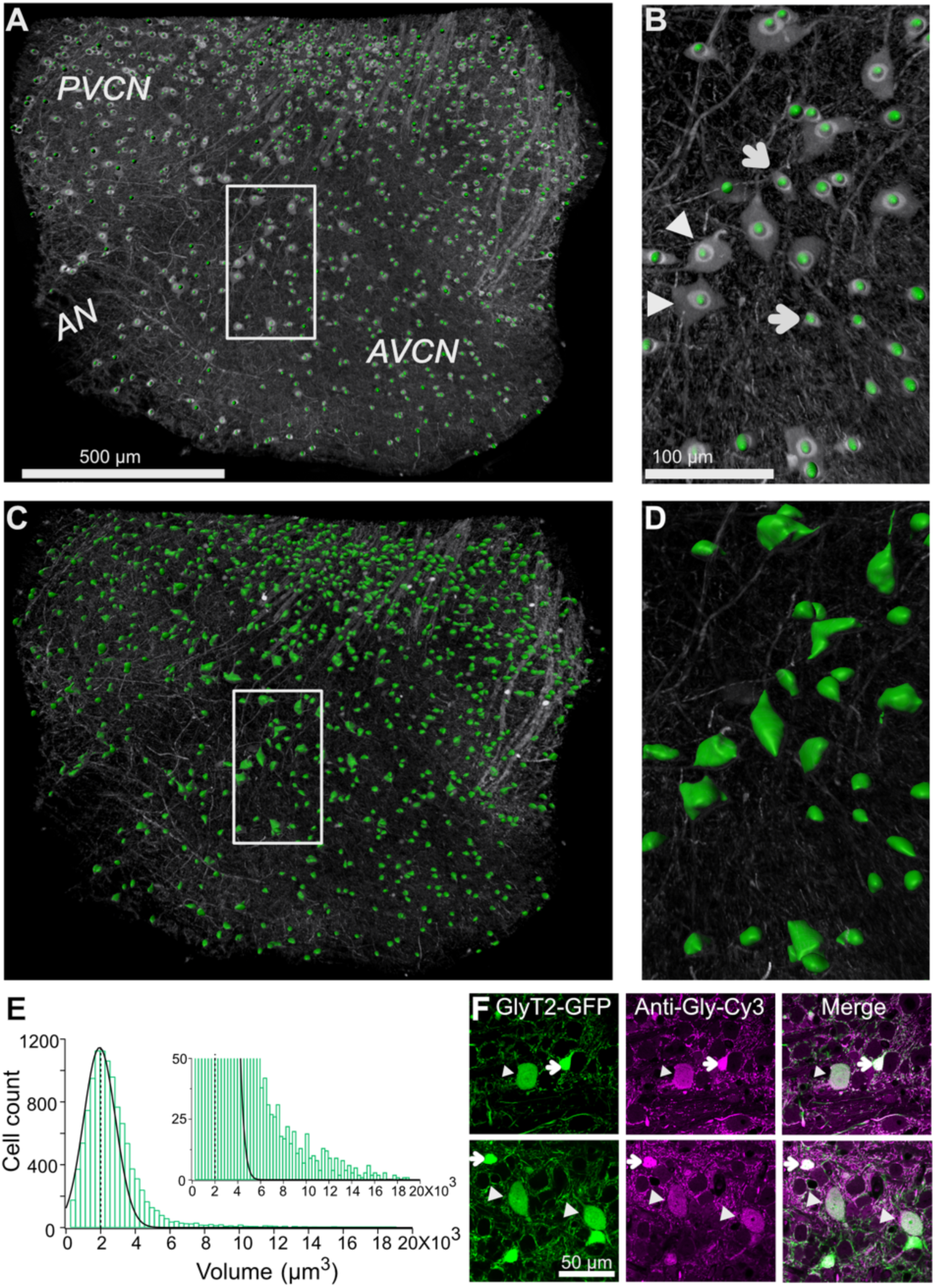
Quantification of glycinergic cells in the VCN. Maximum image projection of a 150 μm thick VCN. Cell count and soma size quantification were restricted to glycinergic cells in the VCN. (**A-B**) Cell counts were quantified using “spot function” in Imaris. Each green dot is counted as one cell. (**B**) There is an anatomically distinct glycinergic cell population in the VCN. Example image shows a mix of large (arrowheads) and small cells (arrows). (**C-D**) Soma volumes were measured with surface rendering program in Imaris. (**E**) Soma size distribution was positively skewed. Dashed line shows the average soma size. The distribution was fitted by Gaussian curve. Majority of the glycinergic cells had small soma size, and minority have large soma size (inset). (**F**) Two representative images of VCN showing large (arrowheads) and small (arrow) glycinergic cells colabeled for glycine. Figure 2-source data 1 Spreadsheet contains all the raw volume data for the 4 CNs used for the quantitative analyses shown in Fig 2. The individual files and their data are present in separate sheets.

To verify that GFP expressing neurons in the VCN from GlyT2-GFP mice were indeed glycinergic, immunohistochemical staining against glycine was performed on VCN sections (Fig 2F). The proportion of GFP expressing cells also positive for glycine was 93.44 ± 1.54%, whereas the proportion of glycine-positive cells also expressing GFP was 99.5% (*n* = 442 cells, *n* = 3 VCNs, 3 mice). Interestingly, RNA probes against the GlyT2 gene (Slc6a5) also clearly reveal both large and small glycinergic cells types in VCN (http://mouse.brainmap.org/experiment/show/69874024).

To study the existence of additional glycinergic sources, we took an intersectional approach to test whether the population of glycinergic neurons in VCN consists of molecularly distinct glycinergic cell types. We used a somatostatin-Cre (Sst-Cre) mouse line that has been used to identify and study somatostatin-containing neurons in the brain. An Sst-Cre::Ai9 mouse line expresses tdTomato in a variety of neurons in the VCN. This mouse line was then crossed to the GlyT2-GFP mouse line. In the resulting cross, large GFP-positive cells were clearly positive for tdTomato; given their size, these are candidate D-stellate cells, and this inference was confirmed by experiments described below on commissural projections.

However, closer examination revealed that the majority of the GFP positive cells were tdTomato-negative and formed a size class distinct from the double-labeled cells presumed to be D-stellate cells. GFP and tdTomato fluorescence was amplified with antibodies to ensure the visualization of cells with lower expression (*n* = 3 VCNs, 2 mice) (Fig 3A, B). There was a strong correlation between GFP and tdTomato expression in presumptive D-stellate cells (Fig 3B, top row). By contrast, smaller cells that were GFP positive were clearly tdTomato negative (Fig 3B, bottom row). Thus, the small glycinergic cells are molecularly distinct from the larger D-stellate cells in that only the latter express cre in this mouse line.

**Figure 3.**
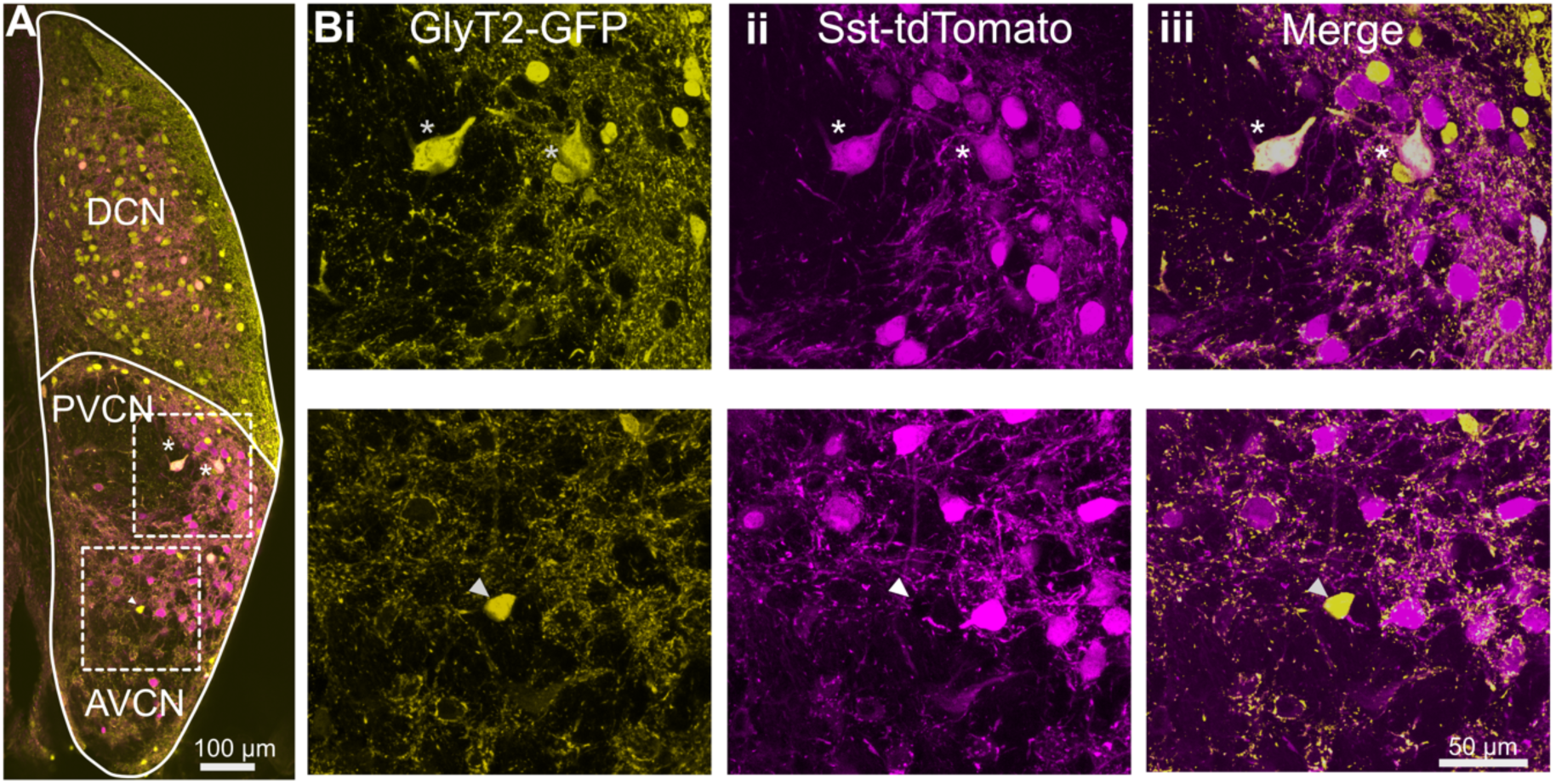
D-stellate cells are molecularly distinct from the small glycinergic cells. (**A**) Coronal section of a CN from an Sst-tdTomato::GlyT2-GFP mouse. (Bi-iii) D-stellate cells show strong colocalization of tdTomato and GFP (inset and *top row*, asterisk) whereas small glycinergic cells were tdTomato negative and GFP positive (*bottom row*).

This conclusion was further probed by examining the distributions of small glycinergic cells and D-stellate cells in the VCN. CNs from Sst-tdTomato::GlyT2-GFP mice were optically cleared the using CUBIC-mount (Fig 4A). We then separately measured the soma volumes of glycinergic cells expressing GFP only and D-stellate cells expressing both GFP and tdTomato using the surface function in Imaris (Fig 4B). We found that double-labeled cells were significantly larger than GFP-only cells (average soma volume, GFP only, 1826.55 ± 12.93 μm^3^ vs. D-stellate cells, 4289.5 ± 98.29 μm^3^, *n* = 2 VCNs, 2 mice). However, numerically, the larger cells types composed only about 12% of all glycinergic neurons in VCN (GFP only, *n* = 3250 ± 55 vs. D-stellate cells, *n* = 380 ± 9) (Fig 4C, inset). Our data thus indicate that D-stellate cell types and small glycinergic cell types are molecularly distinct population of glycinergic interneurons in the VCN, and also show that the small cells compose the vast majority of glycinergic neurons in the VCN.

**Figure 4.**
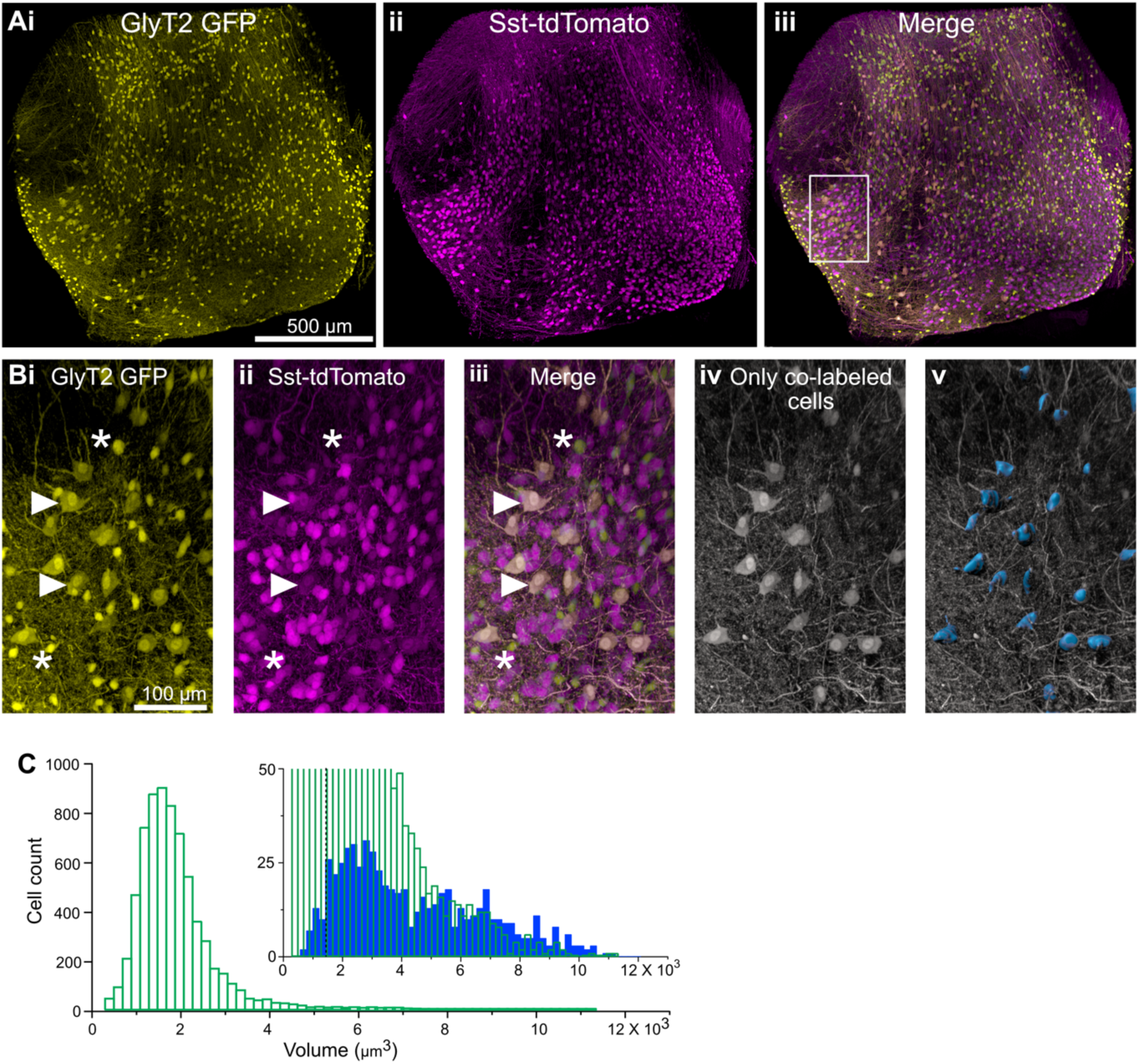
D-stellate cell types are molecularly distinct from the small glycinergic cells. (**A**) 150-μm thick image stacks of optically cleared VCN from Sst-tdTomato::GlyT2-GFP mouse showing GFP (**i**) and tdTomato expressing cells (**ii**). (**B**) An example area from VCN (inset, **Aiii**) showing large GFP (**i**) and tdTomato (**ii**) expressing cells colocalize (**iii**). (**Biv-v**) Only the double-labeled cells are shown and surfaces were rendered to measure to the soma size. (**C**) Soma size distribution of all the glycinergic GFP positive neurons. *Inset*, soma volumes of double-labeled cells. Double-labeled cells were significantly larger than GFP-only cells (average soma volume, GFP only (average soma volume, GFP only, 1826.55 ± 12.93 mm^3^ vs. D-stellate cells, 4289.5 ± 98.29 mm^3^, *n* = 2 VCNs, 2 mice). However, numerically, the larger cells types composed only about 12% of all glycinergic neurons in VCN (GFP only, *n* = 3250 ± 55 vs. D-stellate cells, *n* = 380 ± 9). Figure 4-source data 2 Spreadsheet contains all the raw volume data for the 2 CNs used for the quantitative analyses shown in Fig 4.

### Axon-dendritic arbors of cell types

To further confirm that the small glycinergic cells are a distinct population, we performed individual cell fills by patch clamping GFP-positive neurons with pipettes containing biocytin in the intracellular solution, and reconstructing the filled cells using Neurolucida (Fig 5). Large cells exhibited typical radiate morphology which have been described previously for D-stellate cells in the VCN (Oertel et al., 1990; Campagnola and Manis, 2014; Xie and Manis, 2014) (*n* = 6) (Fig 5A), and henceforth will be referred to as D-stellate cells. Unlike D-stellate cells, the small glycinergic cells had restricted axonal and dendritic arbors (*n* = 17) (Fig 5B). The spread of the axonal-dendritic field was quantified by measuring the longest and shortest axis of the reconstructed structures. We found that small glycinergic cells had significantly restricted branched processes compared with D-stellate cells (longest axis: D-stellate cells, 619.42 ± 45.78 μm vs. small cells, 271.43 ± 26.23 μm, *p* < 0.001, *t*-test; shorter axis: D-stellate cells, 435.62 ± 1.82 μm, *n* = 6 cells *vs*. small cells, 148.60 ± 12.03 μm, *n* = 17, *p* < 0.001, *t*-test) (Fig 5C). We also measured the volume and surface area under the axonal-dendritic field assessed by its convex hull, and found that the field encompassed by the D-stellate cell processes had a larger volume and surface area than that of the small glycinergic cells (Fig 5D, E). Our morphological data indicate that the smaller glycinergic cells are anatomically distinct from the D-stellate cell types. These data provide further evidence that the small glycinergic cells are a distinct cell class of inhibitory neurons in the VCN.

**Figure 5.**
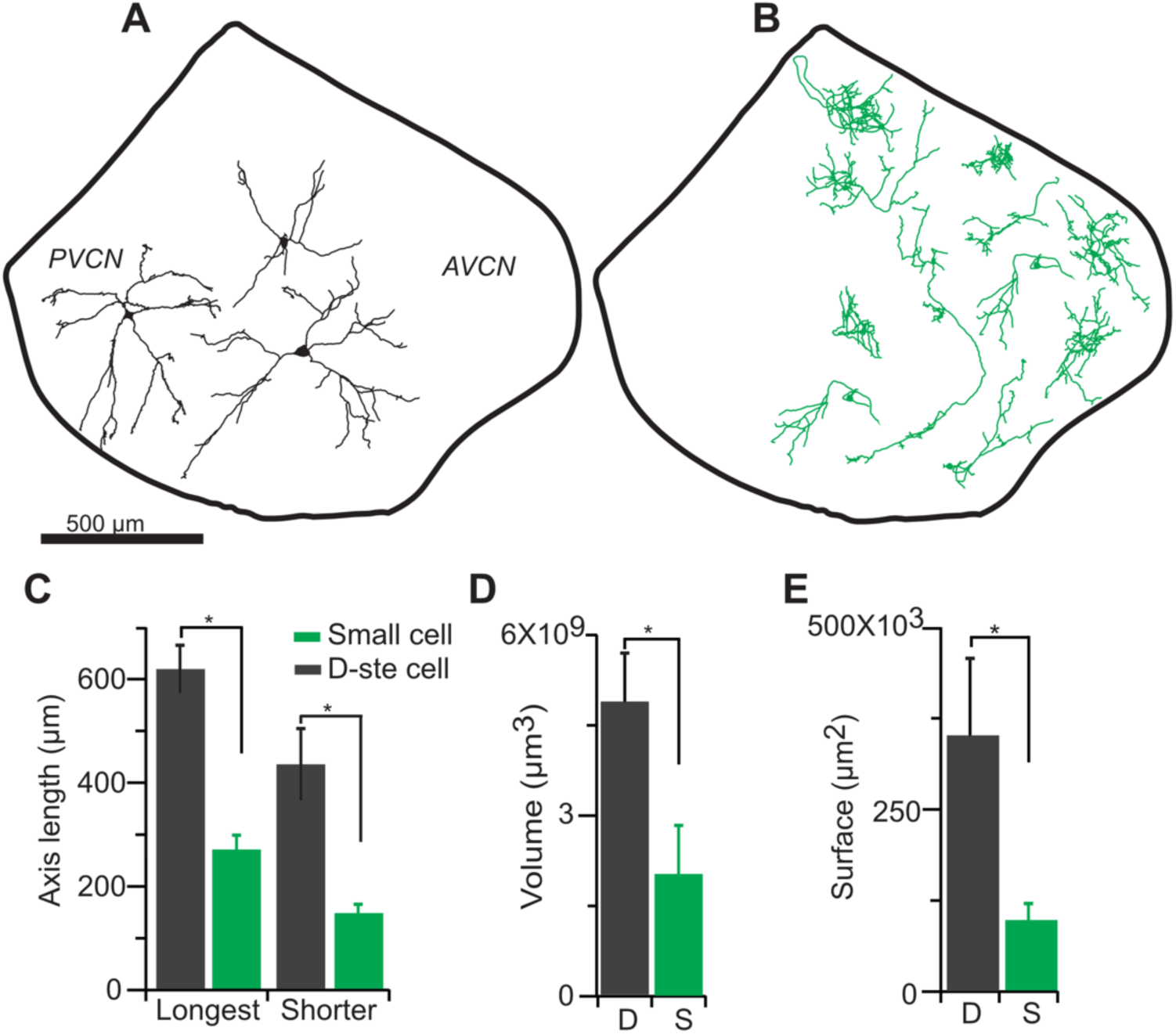
Smaller glycinergic cells are anatomically distinct from D-stellate cells. Biocytin filled neurons were reconstructed using Neurolucida. (**A**) D-stellate cells (large cells) exhibited typical radiate morphology whereas (**B**) the smaller glycinergic cells show restricted axonal and dendritic arbors. (**C**) Spread of axonal-dendritic arbors was quantified by measuring the longest and shortest axis of the reconstructed structures. Smaller glycinergic cells had significantly restricted axonal-dendritic process compared with D-stellate cells (longest axis: D-stellate cells, 619.42 ± 45.78 μm vs. small cells, 271.43 ± 26.23 μm, *p* < 0.001, *t*-test; shorter axis: D-stellate cells, 435.62 ± 1.82 μm, *n* = 6 cells *vs*. small cells, 148.60 ± 12.03 μm, *n* = 17, *p* < 0.001, *t*-test). D-stellate cells had significantly larger volume (volume: D-stellate cells, 4.8 ± 1.2 × 10^9^ mm^3^ *vs*. small cells, 2.02 ± 0.6 × 10^9^ mm^3^, *p* < 0.001, *t*-test) (**D**) and surface area (D-stellate cells, 20.9 ± 1.07 × 10^3^ mm^2^ vs. small cells, 9.8 ± .27 × 10^3^ mm^2^, *p* < 0.01, *t*-test) (**E**) compared with the small glycinergic cells.

### Intrinsic properties

Next, we studied the intrinsic electrical properties of the D-stellate and the small glycinergic cells in the VCN by targeting GFP positive cells from GlyT2-GFP mice for whole-cell recordings. D-stellate cells were identified based on their large soma size and dendrites, visible in patch clamp recordings, while the small cells had obviously smaller somata and less distinct dendrites. Representative responses to depolarizing and hyperpolarizing current injections are shown in Fig 6A,B. All cells showed sustained firing responses at weak-to-moderate depolarizing current steps; however, their responses to strong depolarizing current injections were more diverse (Fig 6A-B, top), particularly as regards the maintenance of spike amplitude during the response. Measurements of membrane properties, action potential shape, and firing properties were collected from all the glycinergic cells (Table 1). We used these three parameters (action potential height, AHP decay and input resistance, Fig 6C, inset) to classify the glycinergic cells into different clusters, in order to probe for distinct inhibitory cell types (Fig 6C). The elbow method for estimating the optimal number of clusters (Fig 6D, inset) indicated 2 or 3 groupings were most likely. We then used K-means cluster analysis to divide the population of all the glycinergic cells into 2 groups (Fig 6C). Cluster analysis based on electrophysiology accurately separated 91.7% (11/12) of visually-identified D-stellate cells (red circles) from the small cells (blue). However, the parameters used to classify the small cells are widely distributed which suggest there may be further subtypes among the small glycinergic cells. When we used a K-value of 3, the ‘small’ cells were further classified into two further groups (Fig 6D, green vs blue circles). The high accuracy of cluster method for classifying the glycinergic cells suggest that small cell types may be solely identified based on the physiological membrane properties of the cells.

**Figure 6.**
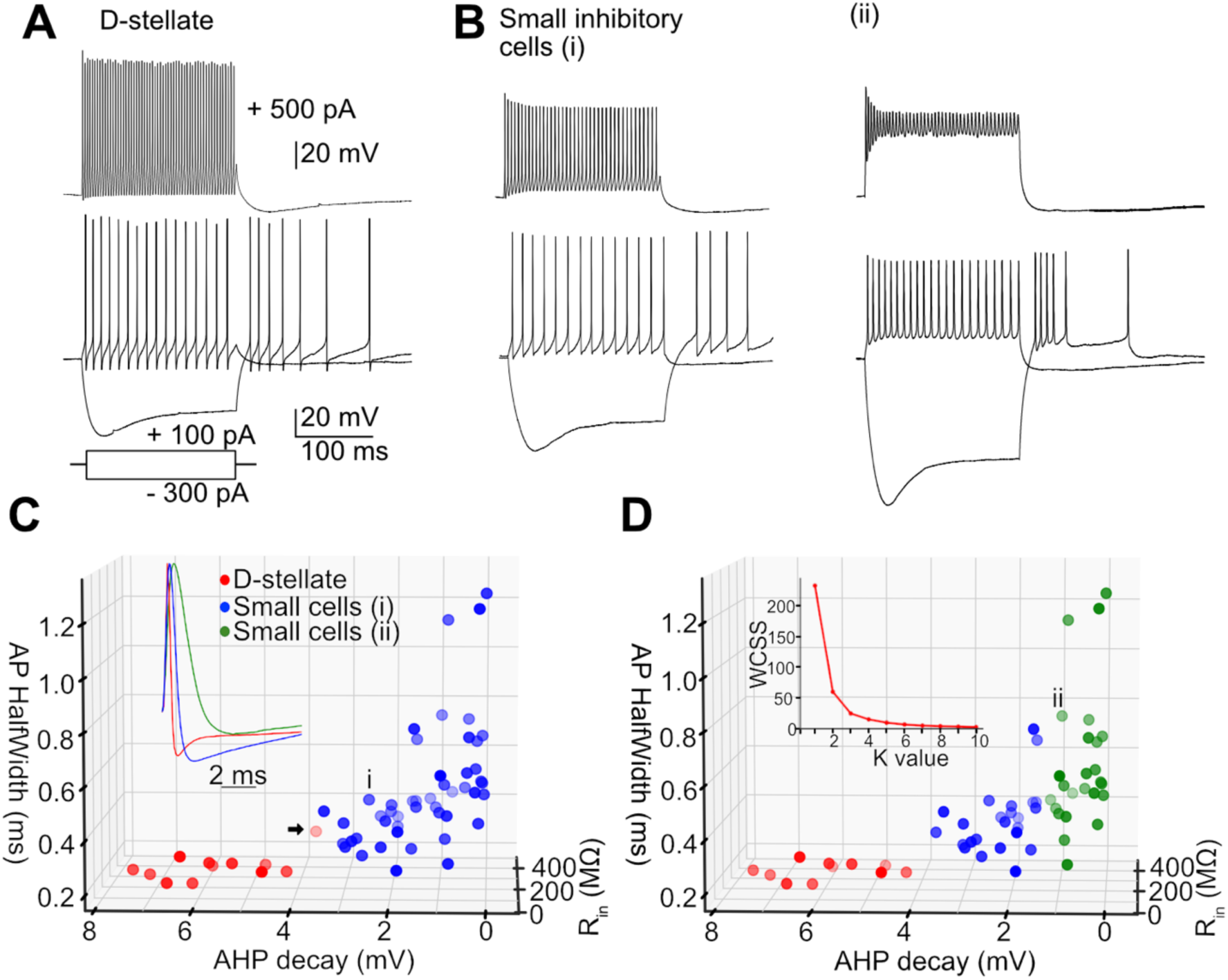
Heterogeneity of spiking properties of the small glycinergic cells. Representative examples of D-stellate cells (**A**) and small glycinergic cell types (**B**) responding to current injections (+500 pA, +100 pA and −300 pA). All cells showed sustained firing responses to +100 pA current injections. Small cells exhibited diverse spike amplitude adaptation in responses to strong (+500 pA) current steps (*top***, Bi-ii**). (**C**) Classification of glycinergic cells based on the action potential height, AHP decay and input resistance. The elbow method was used to calculate the optimal number of clusters (6D, inset (labels, Within cluster sum of squares (WCSS)). Glycinergic cells are group into clusters based on K-means cluster analysis (K=2) (red: D-stellate cells, blue: small cells, color gradient: front (dark) to back (light)). (**D**) K=3 resulted in further classification of small cells into two sub clusters (blue and green). Arrow points to the D-stellate cell included into small cells when K = 3, i and ii represents small cells in blue and green clusters respectively.

**Table 1:** Membrane properties of the glycinergic cells in the VCN. R_in_: Input resistance; AP height: action potential peak potential; AP halfwidth: action potential half width; AP AHP: action potential afterhyperpolarization; AP AHP latency: action potential hyperpolarization latency; Rate of rise of action potential; Threshold: action potential threshold; Overshoot: peak of action potential from 0 mV; Undershoot: Peak of hyperpolarization from baseline; Firing rate: number of spikes/s

| Table 1. Membrane properties of glycinergic cells in the VCN |  |  |
| --- | --- | --- |
| Measure | D-stellate | Small cell |
| R<sub>in</sub> (MΩ) | 114.870 ± 30.7 | 305.540 ± 45.04 |
| AP height (mV) | 70.138 ± 18.75 | 63.923 ± 9.42 |
| AP halfwidth (ms) | 0.310 ± 0.08 | 0.547 ± 0.08 |
| AP AHP (mV) | 4.179 ± 1.11 | 1.546 ± 0.23 |
| AP AHP latency (ms) | 0.875 ± 0.23 | 1.808 ± 0.27 |
| Rate of rise (V/s) | 317.672 ± 84.9 | 288.061 ± 42.47 |
| Threshold (mV) | -45.566 ± 12.17 | -43.462 ± 6.40 |
| Overshoot (mV) | 24.572 ± 6.56 | 20.462 ± 3.01 |
| Undershoot (mV) | -17.788 ± 75 | -14.028 ± 2.06 |
| Firing rate (sp/s) | 213.518 ± 57.06 | 225.720 ± 33.28 |

Many cells in the VCN, such as T-stellate cells (an excitatory principal neuron), exhibit “chopping” responses to acoustic stimuli, such that the peristimulus time histograms of spiking show initial peaks of activity at regular intervals unrelated to the phase of the sound stimulus (Rhode et al., 1983; Young et al., 1988; Blackburn and Sachs, 1989; Smith and Rhode, 1989; Oertel et al., 2011). No definitive *in vivo* recordings of small inhibitory cells in VCN have been reported. To test whether the small cells have the capacity to generate such chopping responses, we recorded from the cells *in vitro* and applied repeated long, constant amplitude depolarizing current steps (200 pA) (Fig 7A). The resulting peristimulus time histograms exhibited features closely resembling choppers *in vivo*, with initial regular peaks and later randomized firing, despite little to no adaptation in firing rate during the whole length of the stimulus (Fig. 7B-D, *n* = 26). This feature raises the possibility that these smaller neurons could give acoustic responses like those of principal cells, and possibly be mistaken for such cells in studies in which no other means for cell identification is used.

**Figure 7.**
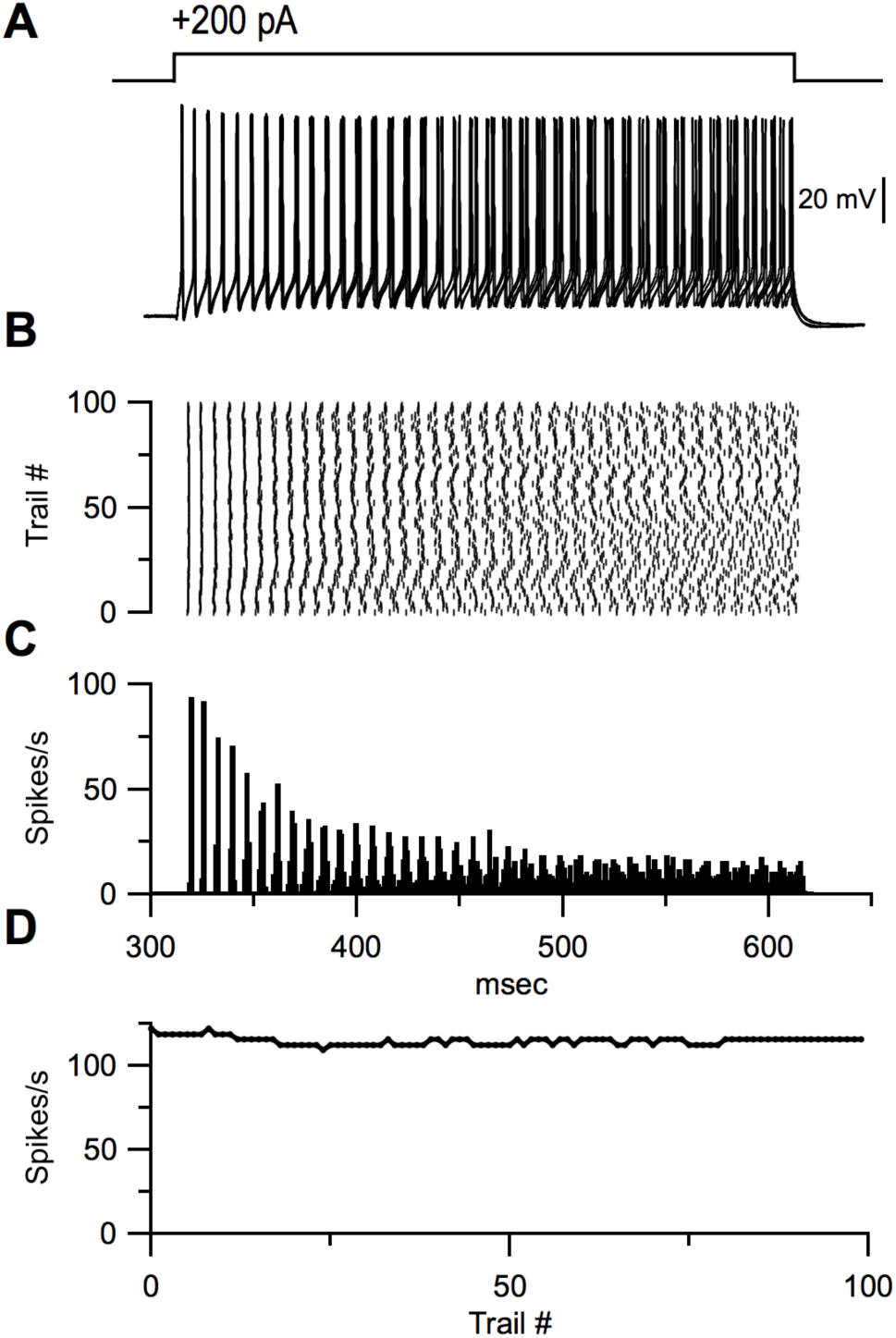
Chopper responses to sustained depolarizing current injections. Representative example of a small glycinergic cell responding to +200 pA current injections (**A**). Raster (**B**) and PSTH (**C**) and firing rate (**D**) of cell in **A**.

### Synaptic connectivity

Auditory nerve fibers (ANFs) are the primary source of excitation to the VCN. However, collaterals of principal cells could in theory provide some local excitation. To test whether the small cells receive ANFs input, we electrically stimulated the auditory nerve root in the presence of strychnine and gabazine to block inhibitory synaptic inputs, and recorded excitatory postsynaptic currents (EPSCs) from small GFP-positive cells. AN root stimulation evoked EPSCs that grew in size with the strength of the shocks (Fig 8A) indicating multiple ANF inputs per postsynaptic cell. Varying stimulus strength showed that each cell receives 3-5 ANF inputs (Fig 8A). The average amplitude of EPSCs evoked with AN root stimulation was 385.7 ± 49.9 pA (*n* = 22). The EPSCs also showed the fast kinetics (τ = 0.80 ± 0.07 ms, half-width = 0.91 ± 0.08 ms, *n* = 22) and had short and consistent synaptic delays of less than 1 ms, consistent with monosynaptic transmission.

**Figure 8.**
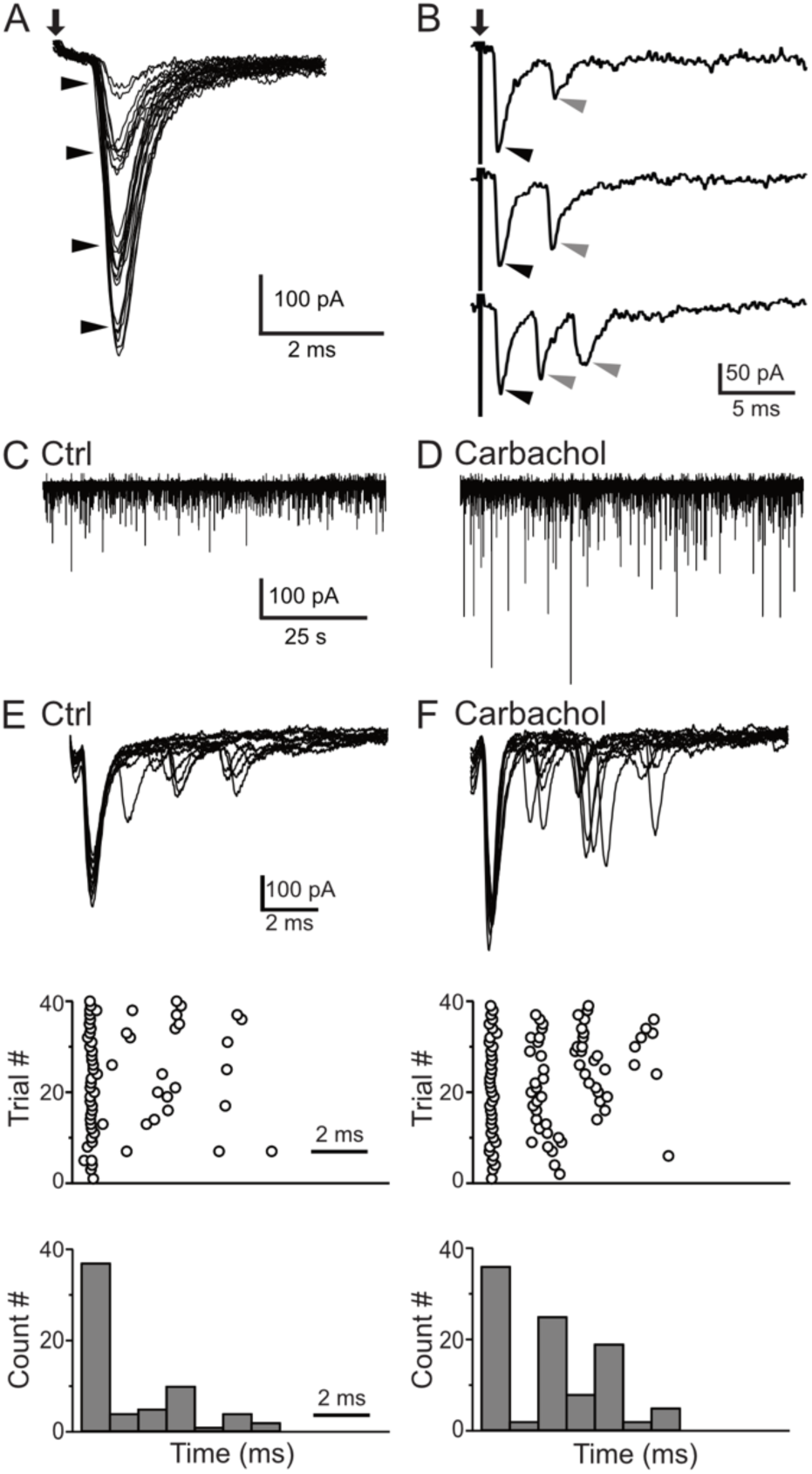
Synaptic inputs to the small glycinergic cells. (**A**) Stimulation of AN root evoked EPSCs with graded amplitudes. The EPSCs amplitude increased with stronger stimulus strength (arrow heads). (**B**) Representative example of disynaptic EPSCs with longer latency in response to AN root stimulation (gray arrow heads). (**C**) Example trace showing sEPSCs measured in small cells under control condition(**C**) and carbachol(**D**). The increased in sEPSCs frequency confirms T-stellate cells as disynaptic source of EPSCs. (**E**) Representative example of eEPSCs in response to AN root stimulation under control conditions (**E**), raster (**G**) and PSTH (**I**) of cells in **E**. (**F-H**) Carbachol application significantly increased the number of delayed EPSCs (short latency events <1 ms, percentage increase = 17 ± 0.11%; longer latency events, percentage increase = 92 ± 0.13%).

In many small cells (45%, 10/22), AN root stimulation evoked EPSCs with distinctly different latencies (Fig 8B). The presence of peaks with different latencies suggests that the small cells also receive polysynaptic excitation from neurons located within the brain slice. Specifically, the first EPSC always had a short synaptic latency consistent with a monosynaptic input (Fig 8B, black arrow heads), but this was often followed by EPSCs two or more ms later (Fig 8B, gray arrow heads). In the VCN, excitatory T-stellate cells has been shown to make local collaterals (Oertel et al., 1990; Ferragamo et al., 1998a) whereas bushy cells appear not to make any local collaterals (Smith et al., 1991; Smith et al., 1993). Moreover, Fujino and Oertel (2001) showed that T-stellate cells can be excited by the cholinergic agonist carbachol. We reasoned that if T-stellate cells are responsible for the delayed EPSCs, the frequency of those EPSCs should be increased if the T-stellate cells were made more excitable with carbachol. To test this idea, carbachol (10 μM) was bath applied while monitoring spontaneous EPSCs (sEPSCs) from small GFP-positive cells. We observed a significant increase in the amplitude and frequency of sEPSCs (Fig 8C, D), as expected if the T-stellate cells were firing spontaneously. In order to test the role of local collaterals in the delayed events, we recorded EPSCs in small GFP-positive cells evoked by AN root stimulation. A representative recording with 10 superimposed traces is shown in Fig 8E. In this cell, we observed a first EPSC with a short latency (< 1 ms) followed by events with longer latencies (> 2 ms). Carbachol (10 μM) was then washed in and EPSCs recorded in response to the same AN root stimulation strength (Fig. 8F). The latencies and numbers of EPSC were quantified and displayed as raster and PSTH plots (1-ms bin; Fig 8 G-J). There was no change in the first EPSC but the events with longer latencies increased in frequency. Across multiple cells (*n* = 5), there was no significant increase in number of events under < 1 ms (percentage increase from control = 17 ± 0.11 %,). By contrast, we observed a significant increase in the number of events with longer latencies (percentage increase from control = 92 ± 0.13 %). These results strongly suggest that the local collaterals from T-stellates are the source of polysynaptic excitatory inputs onto the small glycinergic cells.

We next examined the projections of the small GFP-positive neurons. D-stellate cells project into DCN (Smith and Rhode, 1989; Oertel et al., 2011), as well as to the contralateral CN (Wenthold, 1987; Shore et al., 1992; Schofield and Cant, 1996b; Alibardi, 1998; Doucet et al., 1999; Needham and Paolini, 2003; Doucet and Ryugo, 2006). In order to test whether the small GFP-positive cells also project to the contralateral CN we injected retrogradely-transported fluorescent latex beads (50-100 nl) into the contralateral CN of GlyT2-GFP mice (*n* = 3). Five days after the bead injections, we examined coronal sections of the ipsi- and contralateral CN (Fig 9A, B). Some GFP positive neurons were double-labeled with retrobeads in the ipsilateral CN, however virtually all of these had large somas, consistent with D-stellate cells (length of longest axis = 26.9 ± 3.92 μm, *n* = 47, *n* = 3 CNs) (Fig 9 C). Thus, few if any of the small GFP-positive cells project contralaterally, again indicating that they compose a separate cell class.

**Figure 9.**
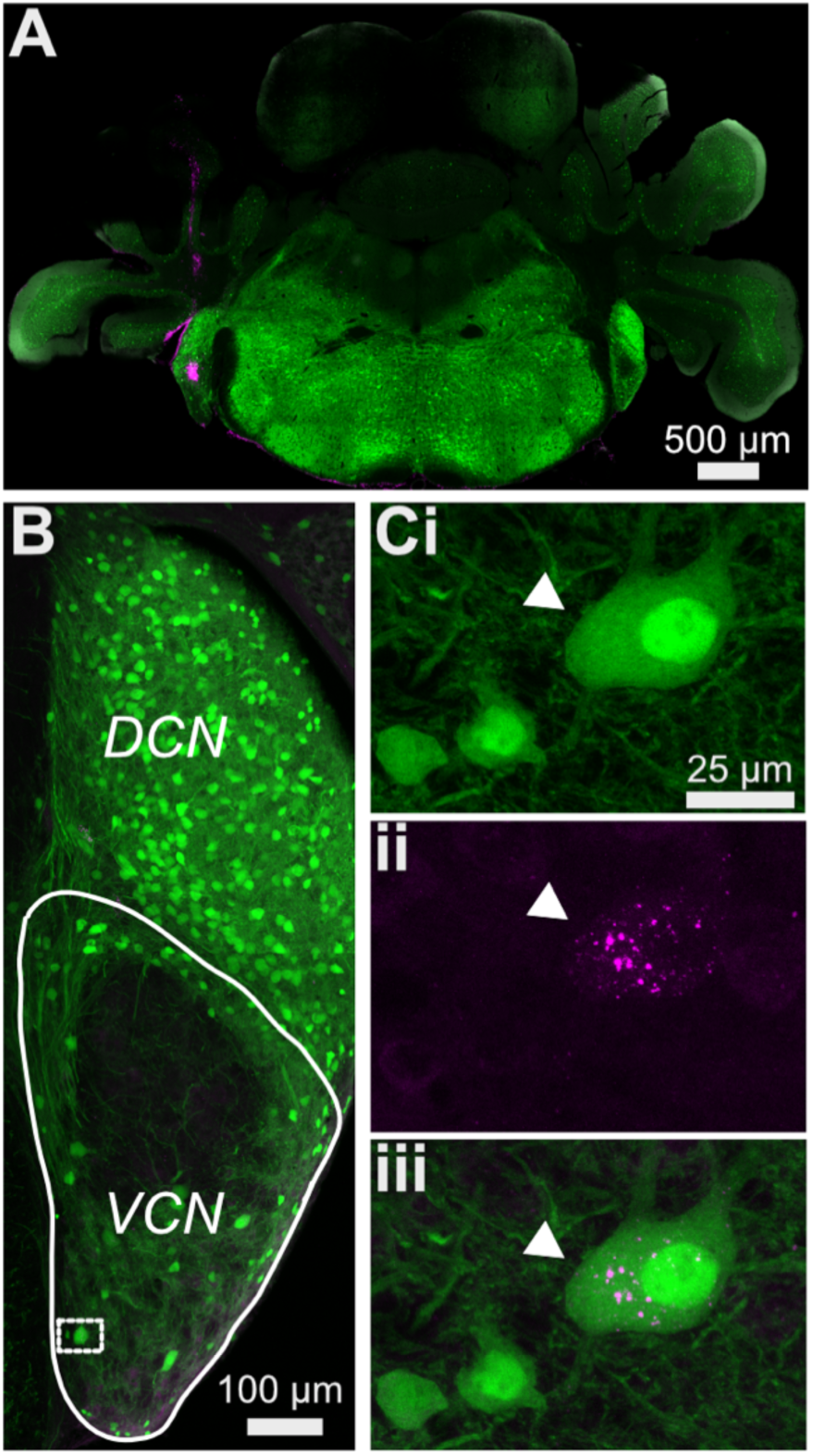
Most small cells do not project contralaterally. (**A**) Retrogradely-transported fluorescent latex beads was injected into the contralateral CN of GlyT2-GFP mice. (**B**) VCN from the Ipsilateral CN was examined for neurons with retrobeads. (**C**) All the double-labeled neurons have large soma size (arrowhead) consistent with D-stellate cells. D-stellate cells (length of longest axis = 26.9 ± 3.92 μm, *n* = 47, *n* = 3 CNs)

Our anatomical reconstruction data (Fig 5) of the small GFP-positive neurons in the VCN suggest these cells might provide local feedforward inhibition to principal cells in the VCN. To test this, we first tried paired-recordings of small cells with principal cells of the VCN, however, out of 40 paired-recordings, we found only a single successful pair. As the yield was very low we used a different approach. Interestingly, we have found that a subset (9/15) of the small GFP-positive cells can be activated by bath application of cholinergic agonists, carbachol (10 μM) (*n* = 15) (Fig 10 A). In contrast, the other major inhibitory sources to VCN including the D-stellate (*n* = 6) (Fig 10 B) and the tubercoluventral cells of the DCN (*n* = 5) (Fig 10 C) failed to respond to carbachol. This lack of carbachol sensitivity in the D-stellate is consistent with (Fujino and Oertel, 2001). To examine whether the small cells provide inhibition to principal cells in the VCN, we enhanced their excitability by bath application of carbachol (10 μM) and recorded inhibitory postsynaptic currents (IPSCs) from bushy cells and T-stellate cells in the presence of excitatory synaptic blockers (10 μM NBQX, 5 μM MK-801). Carbachol application produced a clear increase in the frequency of sIPSCs in both bushy and T-stellate cells (Fig D and E) (inter-event interval, control = 280.4 ± 47.6 ms vs carbachol = 191.2 ± 29.0 ms, *p* < 0.02, *t*-test, *n* = 18) (Fig 10 J). The identity of the principal cells was confirmed using their sIPSCs kinetics. The decay time course of the sIPSCs (bushy cell, τ = 7.22 ± 1.40 ms, *n* = 9, T-stellate, τ = 2.28 ± 0.07 ms, *n* = 9) (Fig 10 F-I), consistent with cell-specific kinetics reported previously (Xie and Manis, 2013b, 2014; Lin and Xie, 2019). Thus, the small GFP-positive, glycinergic cells likely act as local interneurons.

**Figure 10.**
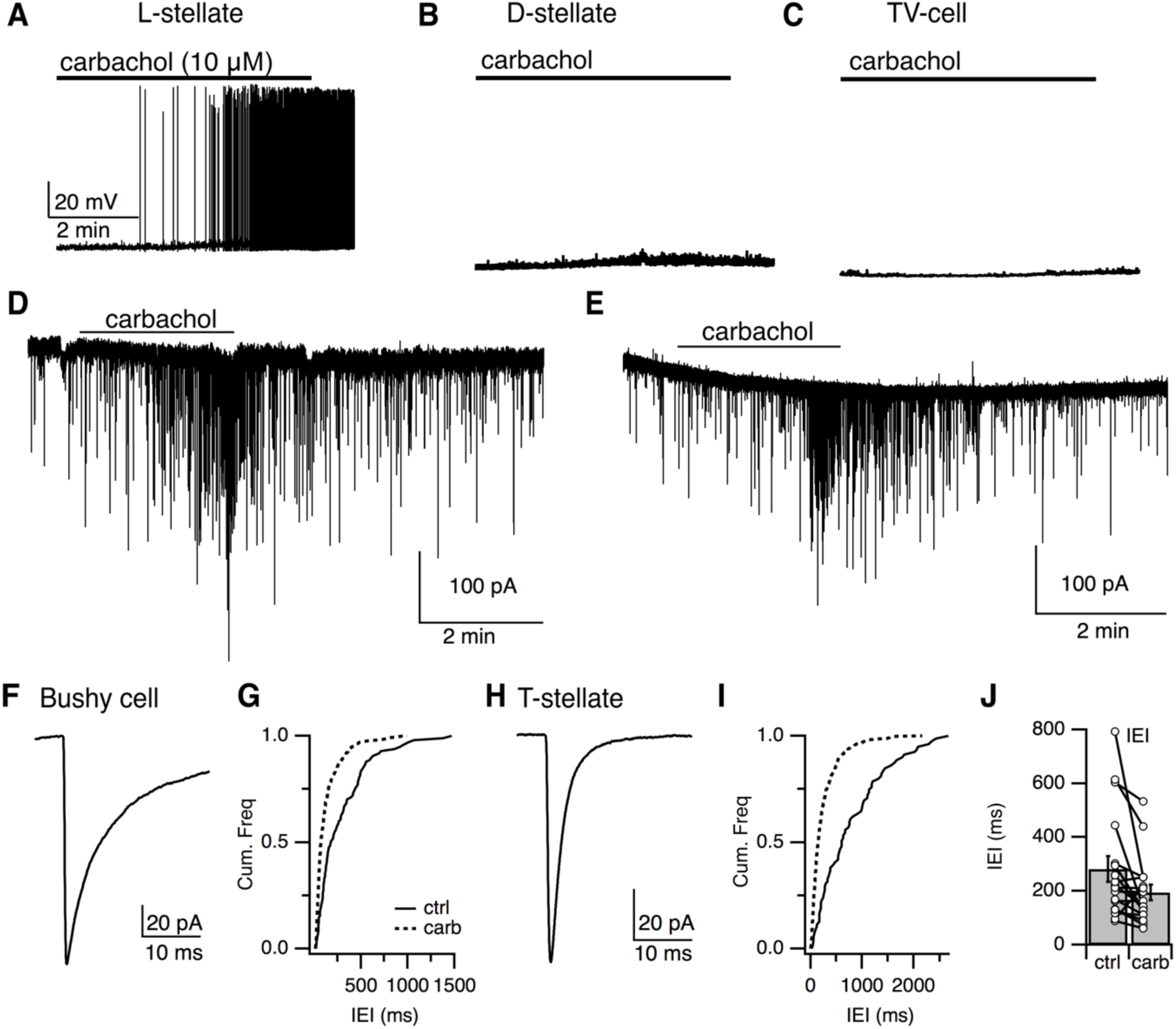
Local stellate cells (L-cells) provide local inhibition to primary cells in the VCN. (**A-C**) Representative responses to cholinergic agonist (carbachol, 10 μM) by small (L-cells) (**A**), D-stellate (**B**), tuberculoventral (TV) cells (**C**). Small cells were activated by carbachol. D-stellate and TV cells failed to respond to carbachol application. (**D**) IPSCs in principal cells, bushy cells (**D**) and T-stellate cells (**E**) in the VCN were induced by carbachol application, confirming that small inhibitory neurons are local interneurons. (**F, H**) Time course of sIPSCs from bushy cells and T-stellate (bushy cell, t = 7.22 ± 1.40 ms, *n* = 9, T-stellate, t = 2.28 ± 0.07 ms, *n* = 9). (**G, I**) Cumulative histogram of inter-event internal under control and carbachol application. (J). Carbachol application resulted in significant increase in sIPSCs frequency (inter-event interval, control = 280.436 ± 47.56 ms vs carbachol = 191.21 ± 28.99 ms, *p* < 0.02, *t-test*)

## Discussion

Subtypes of VCN stellate cells have been named according to the projections of their axons (D-for dorsal, T-for trapezoid bundle). Given their local projections, we term the small, multipolar glycinergic cells ‘L-stellate cells’. This study used transgenic animals, combined with molecular, morphological and electrophysiological techniques to show that the L-stellate cells are molecularly, anatomical, and electrophysiologically distinct from the well-studied, D-stellate cell types. Moreover, the L-stellate cells constitute the vast majority of glycinergic cells in the VCN. They receive monosynaptic auditory nerve inputs and feedforward excitatory inputs from local T-stellate cells, and their axons appear to inhibit principal excitatory cells, the bushy cells and T-stellate cells of the VCN (Fig 11). Therefore, the VCN has distinct glycinergic interneuron populations that likely play distinct roles in auditory processing.

**Figure 11.**
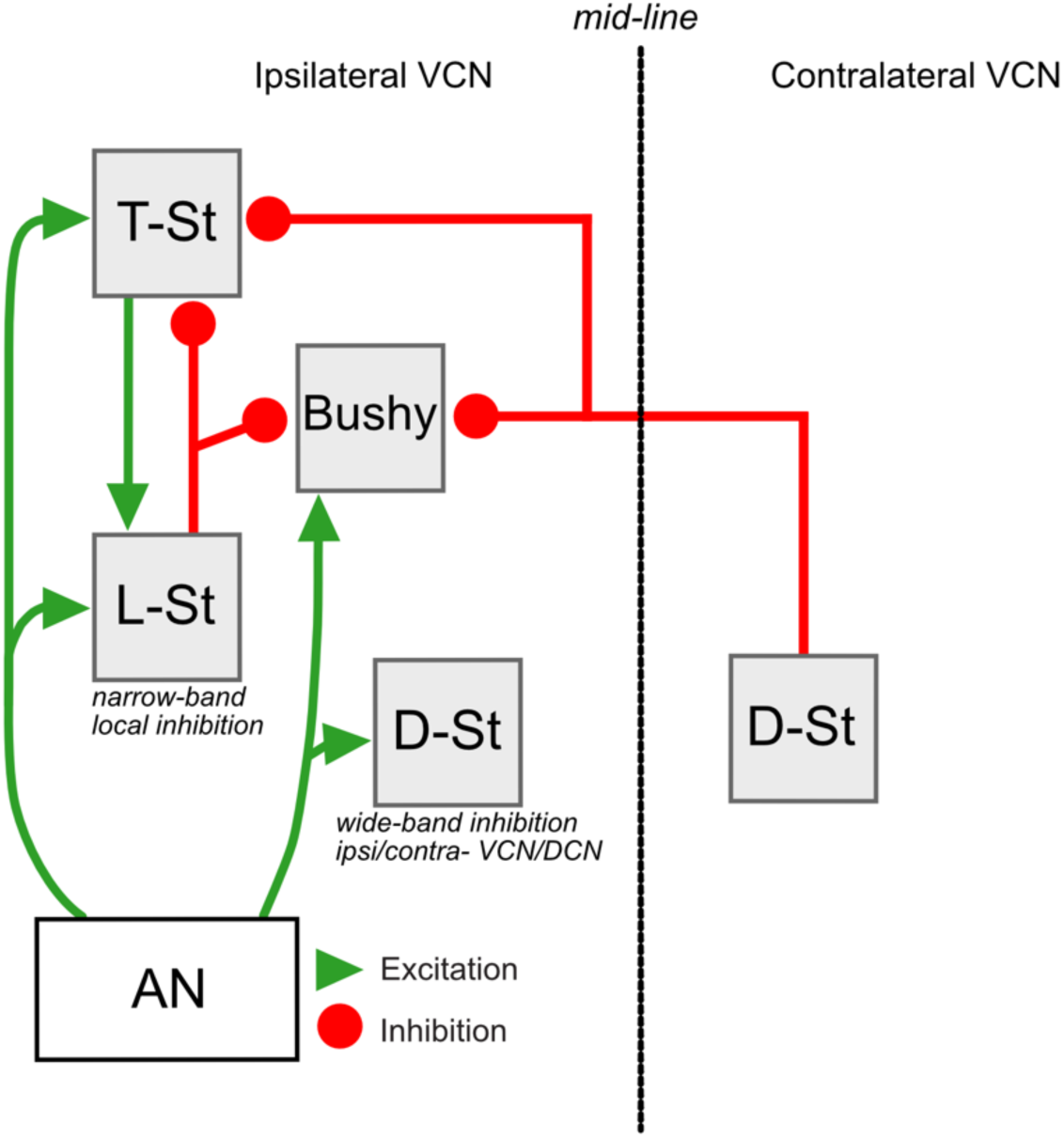
Summary of circuitry in the VCN. Neurons in the ventral cochlear nucleus (VCN) including principal excitatory cells, T-stellate and bushy cells and inhibitory cells, D-stellate and L-stellate cells receive excitatory input from auditory nerve (AN). L-stellate cells also receives polysynaptic inputs from T-stellate cells. L-stellate cells make local inhibitory synaptic contacts on principal cells of the VCN. Principal cells also receive inhibition from D-stellate cells present in the contralateral VCN.

A complex array of inhibitory interneurons sculpt the responses of excitatory neurons and regulate the dynamic circuity across brain regions, and therefore diverse inhibitory cell types would be anticipated to exist in the VCN, the first station of auditory processing. The VCN seemed quite different from this norm in that its only inhibitory cell, the D-stellate cell, is a projection neuron. Although previous studies have hinted at additional small, potentially local glycinergic neurons, it has been a challenge to identify such cells based only on morphological and electrophysiological features. Small cells are generally not targeted well for cell-filling procedures, especially *in vivo*. Electrophysiologically, the glycinergic L-stellate cells show chopper responses, similar to excitatory T-stellate cells (Oertel, 1983; Rhode et al., 1983; Young et al., 1988; Oertel et al., 2011), and thus could have occasionally been mistaken for T-stellate cells.

Morphological and electrophysiological features provide an initial platform for further classification of interneurons. When these approaches are applied to transgenic animals expressing different molecular markers in the inhibitory cell types, a clearer systematic classification system can emerge (Wonders and Anderson, 2006; Cadwell et al., 2016; Fuzik et al., 2016; Tremblay et al., 2016; Pelkey et al., 2017). In the Sst-tdTomato::GlyT2-GFP line, large D-stellate cell types express both GFP and tdTomato, whereas smaller L-stellate cell types were only GFP-positive.

Although we have identified two major glycinergic interneuron populations, further studies may reveal additional diversity within the groups described in this paper. Our anatomical data show a variety of somatic, dendritic, and axonal morphologies within the L-stellate cell types. Electrophysiologically, L-stellate cell types display heterogeneity in their membrane properties, likely a consequence of diverse ion channel expression. We also found that only a subset of L-stellate cells were sensitive to the cholinergic agonist, carbachol. Previous studies have reported that medial olivocochlear axons make synaptic contacts with T-stellate cells and small unidentified cells in the VCN (Benson and Brown, 1990; Benson et al., 1996; Fujino and Oertel, 2001). As some L-stellate cells were activated by bath application of carbachol, this suggests that the L-stellate cells might also receive inputs from the medial olivocochlear pathway. Single-cell transcriptomic analysis will likely allow further definition of subpopulations of glycinergic interneurons in the VCN (Jiang et al., 2015; Cadwell et al., 2016; Fuzik et al., 2016; Shrestha et al., 2018; Sun et al., 2018).

### Local inhibitory inputs in the VCN

In comparison to D-stellate cells, L-stellate cells have smaller soma sizes and exhibit tightly branched axons and dendrites. These restricted dendritic and axonal processes suggest that L-stellate cells have narrow frequency-band receptive fields. This morphology is very different from the D-stellate cells, which has been termed the “wide-band inhibitor” (Nelken and Young, 1994; Arnott et al., 2004). The potential for narrow band excitation of the L-stellate cells could provide a means for lateral inhibition to excitatory cells in nearby isofrequency lamina in VCN. Indeed, when we activated the smaller L-stellate cells with carbachol, we recorded IPSCs from both bushy and T-stellate cells. This provides evidence that L-stellate cells could in principal mediate lateral or side-band inhibition. Lateral inhibition has been proposed to improve tuning properties and temporal precision, and stabilizes neuronal firing responses to acoustic inputs (Gai and Carney, 2008; Keine and Rubsamen, 2015). Indeed, as the D-stellate cells are numerically a minority of the glycinergic cell populations compared to the L-stellate cells, we proposed that the majority of local inhibitory inputs onto excitatory neurons could originate from L-stellate cells.

D-stellate cells sends their axons to the DCN and also project to contralateral VCN via the commissural pathway (Schofield and Cant, 1996a; Doucet and Ryugo, 2006). The commissural D-stellate cell connects ipsi- and contralateral CN. Previous tract tracing studies in rat reported a presence of small commissural glycinergic cells (Doucet and Ryugo, 2006), however we did not observe any L-stellate cells retrogradely labelled in our mouse study, consistent with the restricted axonal morphology of the L-stellate cells. However, our cell-filling experiments only sampled a subset of cells, and it remains possible that some L-stellate cells do project outside the VCN, perhaps to the dorsal cochlear nucleus or granule cell region. Indeed, many L-stellate cells that were located near the margins of VCN, near granule cell regions. These could be the small multipolar marginal cells described by Doucet and Ryugo (2006). However, inhibitory Golgi cells are located in and near the granule cell layer (Mugnaini et al., 1980; Ferragamo et al., 1998b; Irie et al., 2006; Yaeger and Trussell, 2015), so it is possible that some L-stellate cells were Golgi cells.

In addition to local glycinergic sources, VCN receives projecting glycinergic inputs from the tuberculoventral cells of the DCN (Wickesberg and Oertel, 1988, 1990; Wickesberg et al., 1991; Xie and Manis, 2013b; Campagnola and Manis, 2014). Tuberculoventral cells have narrow frequency-receptive fields and project to a tonotopically matched region in the VCN. Therefore, tuberculoventral cells have been proposed to provide narrow band inhibition, but at present it is unclear whether the local glycinergic inputs from L-stellate cells and the long-range inputs from tuberculoventral cells play different roles in sculpting the responses of the excitatory neurons. Conceivably, VCN principal cells could receive input from both subtypes of interneuron, and the differential activation and modulation of those interneurons could aid in fine tuning auditory processing under different conditions.

## Materials and Methods

### Animals

All procedures were approved by the Oregon Health and Science University’s Institutional Animal Care and Use Committee. GlyT2-GFP mice (Zeilhofer et al., 2005) and Somatostatin(Sst)-Cre tdTomato::GlyT2-GFP of either sex, postnatal days(P) 17-30 were used in this study. GlyT2-GFP mice were backcrossed into the C57BL/6J and maintained as heterozygous (Roberts et al., 2008). Sst-tdTomato::GlyT2-GFP mice were generated as follows: Homozygous Sst-IRES-Cre knock-in mice (Jackson Laboratory) were crossed with homozygous Ai9 (RCL-tdTomato) reporter mice (Jackson Laboratory), enabling Sst-IRES-Cre-dependent expression of tdTomato. Offspring from the F1 generation (Sst-IRES-Cre+/-, Ai9(RCL-tdTomato)+/-) were crossed with homozygous GlyT2-GFP mice, whose offspring will be referred to as Sst-tdTomato::GlyT2-GFP. At 5 days postnatal, a fluorescent stereoscope (Leica Microsystems) was used to identify transgenic mice positive for either tdTomato or GFP.

### Brain-slice preparation

Animals were anesthetized with isoflurane and decapitated. The brain was quickly removed and placed into ice-cold sucrose cutting solution. Sucrose solution contained (in mM) 76 NaCl, 26 NaHCO_3_, 75 sucrose, 1.25 NaH_2_PO_4_, 2.5 KCl, 25 glucose, 7 MgCl_2_ and 0.5 CaCl_2_, bubbled with 95% O_2_: 5% CO_2_ (pH 7.8, 305 mOsm). Parasagittal slices of cochlear nucleus were cut at a slight angle from sagittal, to best preserve a straight projection of the auditory nerve (Ngodup et al., 2015). The slices of cochlear nucleus were cut at 220-300 μm in ice-cold sucrose solution on a vibratome (Leica VT1200S or Campden 7000smz-2). Slices were transferred into standard artificial cerebrospinal fluid (ACSF) containing (in mM) 125 NaCl, 26 NaHCO_3_, 1.25 NaH_2_PO_4_, 2.5 KCl, 20 glucose, 1 MgCl_2_, 1.5 CaCl_2_, 2 Na-pyruvate, and 0.4 Na L-ascorbate, bubbled with 95% O_2_:5% CO_2_ (pH 7.4, 300-310 mOsm). Slices recovered at 34 °C for 40 min and maintained at room temperature until recording.

### Electrophysiology

Slices were transferred to a recording chamber and perfused with standard ACSF (∼34 °C). Cells were viewed using an Olympus BX51WI microscope with a 60X objective, equipped with an infrared Dodt contrast, CCD camera (Ret 2000R, QImaging) and fluorescence optics. In slices from GlyT2-GFP, glycinergic cells in the ventral cochlear nucleus (VCN) were identified by their GFP expression. Recording pipettes were pulled from 1.5-mm OD, 0.84-mm ID borosilicate glass (WPI-1B150-F) to a resistance of 2–4 MΩ using a horizontal puller (Sutter Instrument P97). The internal recording solution contained in (mM) 113 K gluconate, 2.75 MgCl_2_, 1.75 MgSO_4_, 0.1 EGTA, 14 Tris_2_-phosphocreatine, 4 Na_2_-ATP, 0.3 Tris-GTP, 9 HEPES with pH adjusted to 7.25 with KOH, mOsm adjusted to 290 with sucrose (ECl −84 mV). For a few voltage-clamp experiments, we used another internal solution containing (in mM) 5 CsCl, 1 MgCl_2_, 4 Mg-ATP, 0.4 Tris-GTP, 5 EGTA, 14 Tris_2_-phosphocreatine, 4 Na_2_-ATP, 10 HEPES, 3 QX-314 (pH 7.2, 290 mOsm). Whole cell patch-clamp recordings were made using a Multiclamp 700B amplifier and pCLAMP 10 software (Molecular Devices). Signals were digitized at 20-40 kHz and filtered at 10 kHz by Digidata 1440A (Molecular Devices). In voltage clamp, cells were held at −70 mV, with access resistance 5‒20 MΩ compensated to 50-60%. In current clamp, resting potential was maintained at −60 to −70 mV. To isolate excitatory postsynaptic currents, inhibitory synaptic blockers SR-95531 (10 µM) and strychnine (0.5 µM) were added to the bath solution whereas to isolate inhibitory currents, excitatory synaptic blockers, NBQX (10 µM) and MK-801 (5 µM) were added to the bath solution. The auditory nerve root was stimulated with brief voltage pulses (100 µs) using a stimulus isolation unit (Iso-Flex, A.M.P.I) via a bipolar microelectrode placed in the nerve root.

### Morphology

Morphological studies of individual cells were made by including 0.2‒0.4% biocytin (B1592, Molecular Probes) in the recording pipette solution. After loading each cell for 20 minutes, the recording electrode was slowly retracted, and the slice fixed overnight in 4% (w/v) buffered paraformaldehyde. After fixation, slices were rinsed in PBS and stored for up to a week at 4 °C in PBS until processing for biocytin labeling. To visualize biocytin labeling, slices were permeabilized in 0.2% Triton X-100 solution (in PBS) for 2 h at room temperature. Slices were incubated 0.3% H_2_O_2_ for 30 min to quench endogenous peroxidase, rinsed with PBS, incubated in ABC reagent (Vector Laboratories) for 2 h, rinsed with PBS, and then incubated for 3‒4 min in diaminobenzidine (DAB) solution containing 0.05 M Tris buffer, 10 mg/ml nickel ammonium sulfate, 50 mM imidazole, 1 mg (1 mg/100 μl) DAB, and 0.3% H_2_O_2_. Slices were then rinsed, mounted on glass slides, dehydrated in an ascending series of alcohols, delipidized in xylene, and cover slipped with Permount. DAB-stained cells were visualized with a Zeiss Axio Imager M2 using a 40X oil immersion objective and reconstructed using Neurolucida (MBF Bioscience). Analyses of dendritic and axonal processes were made in Neurolucida explorer.

### Immunohistochemistry

For glycine labeling, mice were deeply anesthetized with isoflurane and perfused transcardially with 0.9% warm saline followed by a fixative containing 2% glutaraldehyde and 1% paraformaldehyde buffered in PBS. Brains were removed and post-fixed for 30‒60 minutes. Sagittal sections were cut at 50-µm thickness on a vibratome (Leica VT1000S). Slices were rinsed in PBS and then incubated in fresh 1% NaBH_4_ for 30 minutes to reduce autofluorescence from glutaraldehyde fixation. Slices were rinsed extensively in PBS.

For colabeling studies of Sst-tdTomato and GlyT2-GFP-positive neurons, Sst-tdTomato::GlyT2-GFP mice were deeply anesthetized with isoflurane and perfused transcardially with 0.9% warm saline followed by 4% paraformaldehyde buffered in PBS. Brains were removed and post-fixed overnight at 4 °C. Sections were cut at 50-µm thickness on a vibratome (Leica VT1000S).

All the sections were then blocked in 2% bovine serum albumin (BSA), 2% fish gelatin and permeabilized in 0.2% Triton X-100 in PBS for 2 h at room temperature on a shaker table. Next, the sections were incubated in primary antibody solution containing: primary antibody, 2% BSA, 2% fish gelatin PBS with 0.2% Triton X-100 for 24‒48 h at 4 °C. The sections were then washed three times, 10-mins each, in PBS, and incubated in secondary antibody solution for 2 h at room temperature or 24‒48 h at 4 °C on a shaker table. The following primary antibodies were used: 1:500 anti-Glycine (Ab139, Abcam), 1:2,000 chicken anti-GFP (GFP-1020, Aves Labs) and 1:500 rabbit anti-DsRed (632496, Clontech). The following secondary antibodies were used: 1:500 Cy3-conjugated donkey anti-rabbit antibody (Jackson Immuno Research), 1:500 donkey anti-chicken conjugated to Alexa Fluor 488 (703-545-155, Jackson Immuno Research) and 1:400 donkey anti-rabbit conjugated to Alexa Fluor 594 (711-585-152, Jackson Immuno Research). Sections were rinsed in PBS three times, 10 minutes for each rinse, and in a few cases sections were subsequently incubated in 4% paraformaldehyde in PBS for one hour. Sections were then mounted on microscope slides and cover slipped with Fluoromount-G (Southern Biotech). Images were acquired using a Zeiss LSM 780 confocal microscope.

### Thick tissue clearing

Mice were deeply anesthetized with isoflurane, then perfused transcardially with 0.9% warm saline followed by 4% paraformaldehyde buffered in PBS. Brains were removed and post-fixed overnight at 4 °C. The entire cochlear nucleus (450-500 μm) was cut from the brainstem using a vibratome (Leica VT1000S). Because the cochlear nucleus sits at the lateral edge of the brainstem, only a single cut was required to prepare the specimen. Samples were washed in 0.1 M PBS before clearing for 72 h at room temperature on a shaker table with an accelerated ‘clear unobstructed brain imaging cocktail’ (CUBIC)-mount solution (Lee et al., 2016) containing sucrose (50 %, w/v), urea (25%, w/v), and N,N,N′,N′-tetrakis (2-hydroxypropyl)ethylenediamine (25%, w/v) dissolved in 30 ml of dH_2_O. Once each sample was cleared and almost transparent, it was then mounted on a glass slide with 0.5 mm deep silicone spacers (EMS) in CUBIC-mount. The images were acquired within 24 h to reduce volume changes in CUBIC-mount (10-20% increase in volume after incubation). The samples were imaged using a fast Airyscan LSM880 microscope with a 25X objective. Tiles and z-stack of whole cochlear nucleus were imaged at 5-μm step size with the exception of one sample where images were obtained at 2-μm step size. Images were post-processed and individual tiles were stitched together using Zeiss Zen Black software. Analysis of cell count and soma volume were quantified using Imaris software (Bitplane 12.1). For optimal processing of large file sizes, files were separated into 3-4 smaller files each containing (30 GB) ∼150 μm of cochlear nucleus for faster processing. The image stacks were further processed for background corrected and normalization in Imaris. VCN was selected for quantification by drawing regions of interest around the VCN followed by applying a mask outside the region of interest. The dorsal border of the VCN was defined by the granule cell lamina, and the medial border by the absence of glycinergic somata (Muniak et al., 2013). For cell count quantification, we used the built-in “spot detection” algorithm in Imaris in which the program places a “spot” on the soma of each cell. The spots are used for counting of GFP positive cells in the VCN. Automatically detected spots were verified and corrected manually. Thus, if the computer failed to detect a cell or erroneously placed a spot on a background bright spot, we manually added spots and deleted spots respectively. To test the validity of automated spot function, certain regions of VCN were manually counted and compared against the spot function. The spot function detected more than 95% of all cells present in that area. For volume measurements, we used the in-built “surface” rendering function in Imaris. The program rendered 3D surfaces on the soma of GFP positive cells. Rendered surfaces were used to extract the volume statistics. Surfaces were also verified and corrected manually.

### Colocalization Assay

For colocalization analysis of tdTomato with GFP in the VCN, images were obtained from CUBIC-cleared VCN of Sst-tdTomato::GlyT2-GFP transgenic mice. Images were acquired at 5-μm steps with fast Airyscan LSM880 as described above. Images were post-processed and stitched using Zeiss Zen Black software. Images were analyzed using the “ImarisColoc” function in Imaris. 150-μm thickness of VCN were processed separately. ImarisColoc function allows to process the overlap between the two-color channels in an image. Minimum threshold was selected for the two-color channels. A new channel was generated that only contained colocalized voxels. Next, we used the surface rendering program, which turns voxels into solid objects, which were used to measure the volumes on double-labeled cells.

### Retrobead injections

GlyT2-GFP mice were anesthetized with isoflurane and placed in a stereotaxic frame (David Kopf). Animal temperature was maintained near 37 ° C with a heating pad (T/pump Gaymar). The scalp was reflected, a portion of skull above the left cerebellum was opened. VCN was located by stereotactic coordinates starting from the surface junction point of the inferior colliculus, cerebellar lobule IV-V and simple lobule (9.5 mm lateral, 7 mm rostral, 4 mm depth). Glass capillaries (Wire Trol II, Drummond Scientific) were pulled on a horizontal puller (P-97, Sutter) and then beveled using a diamond lapping disc (0.5 μm grit, 3M DLF4XN_56611X) to an inside diameter of 20‒30 μm (Balmer and Trussell, 2019). Glass capillaries were advanced into the VCN with a microdrive (IVM-500, Scientifica) at a rate of 10 μm/s. 50‒100 nl of red retrobeads (LumaFluor Inc.) were injected using a single axis manipulator (MO-10, Narishige) and pipette vice (Ronal). 5-min waiting periods were allowed before and after injections. The skin was sutured and mice were allowed to recover for 5‒6 days. After recovery, mice were deeply anesthetized with isoflurane and then perfused transcardially with 0.9% saline followed by 4% paraformaldehyde buffered in PBS. Brains were removed and post-fixed overnight at 4 °C. Coronal sections were cut at 50-µm thickness on a vibratome (Leica VT1000S). Confocal images of the sections were acquired using LSM 780. Images were analyzed using ImageJ.

### Pharmacology

All drugs in the slice experiments were bath applied. Receptor antagonists used this study included: NBQX (AMPA receptors: Sigma), MK-801 (NMDA receptors; Sigma), SR-95531 (GABA_A_R; Tocris), strychnine (glycine receptors; Sigma). Acetylcholine receptors were activated using the non-selective cholinergic agonist, carbamoylcholine chloride (carbachol) (Tocris).

### Experimental design and statistical analyses

Electrophysiological data were analyzed using pClamp 10.4 software (Molecular Devices), Axograph, or IGOR Pro v6.3 or v8 (WaveMetrics). Figures were made using IGOR Pro, Affinity Designer and Adobe Illustrator. Statistics were performed in IGOR Pro, Axograph, Python, Microsoft Excel or Prism (GraphPad). For statistical analysis, groups were compared with paired or unpaired *t*-test. Cluster analysis were performed using sklearn.cluster.KMeans in Python and figures were made using matplotlib.pyplot. Error bars are represented as mean ± SEM unless otherwise stated.

## Acknowledgements

We thank Ben Suter, Taro Kiritani and Carl Peterson for permission and assistance with the CUBIC protocol, Stefanie Kaech Petrie, Aurelie Snyder, Crystal Chaw and Brian Jenkins from the Advanced light Microscopy Core, The Jungers Center for assistance with microscopy, T. Balmer, T. Garett, H. Hong, L. Moore, J. Tang, D. Zeppenfeld for helpful comments on the manuscripts. We also want to thank M. J. Murdock Charitable Trust for endowing funds for microscopes at the OHSU Microscopy Core. These experiments were supported by Hearing Health Foundation’s Emerging Research Grant to TN, National Institute of Health (NIH) Grants R01 NS028901 and DC004450 to LOT, NINDS P30NS061800 to Imaging Center, OHSU. Gabriel E. Romero is a Howard Hughes Medical Institute Gilliam Fellow.

